# Tetramer formation of CpoS facilitates Inc-Inc interactions during *Chlamydia trachomatis* infection

**DOI:** 10.1101/2024.12.01.621710

**Authors:** Xavier Tijerina, C.A. Jabeena, Robert Faris, Zhen Xu, Parker Smith, Nicholas J. Schnicker, Mary M. Weber

## Abstract

*Chlamydia trachomatis* (*C.t.*), the leading bacterial cause of sexually transmitted infections, replicates within a unique intracellular compartment called the inclusion, which is modified by secreted proteins known as inclusion membrane (Inc) proteins. Here we further characterize CpoS, an Inc previously shown to be critical for replication and inclusion development. We demonstrate that CpoS directly binds multiple coiled-coil domain-containing Incs while simultaneously engaging Rab GTPases at a separate site. Notably, CpoS-InaC interactions facilitate the recruitment of select Arfs to the inclusion membrane, while Rab recruitment occurs independtly of these interactions. Biochemical and biophysical analyses revealed that Incs self-oligomerize, forming higher-ordered structures, with CpoS adpoting a tetrameric structure resembling eukaryotic SNAREs. We propose these assemblies likely serve as scaffolds to orchestrate vesicle docking, tethering, and fusion. Our findings underscore the intricate interplay between bacterial and host factors, revealing that *C.t.* leverages both Inc-Inc interactions and host protein engagement to manipulate vesicular trafficking and sustain infection.

## Introduction

*Chlamydia trachomatis* (*C.t.*) causes the most common bacterial sexually transmitted infection and is the leading cause of infectious blindness worldwide. Genital infections caused by *C.t.* are often asymptomatic, leading to prolonged infections with severe complications such as pelvic inflammatory disease, infertility, ectopic pregnancy, and an increased risk of cervical and ovarian cancer^1–4^. Although these infections can be treated with antibiotics, reinfection is common due to a lack of long-term immunity and the absence of a vaccine^5–7^. A deeper understanding of the cellular and molecular mechanisms that *C.t.* employs to cause disease is crucial for developing improved therapeutics and guiding vaccine development.

Chlamydial replication requires formation of a privileged niche - termed the inclusion-which is extensively modified early in infection through the incorporation of unique type III secreted effector proteins called inclusion membrane proteins (Incs)^8,9^. Thirty seven Inc proteins have been confirmed to localize to the inclusion membrane during infection, representing 5% of *C.t.*’s highly reduced genome^10^. Given their strategic positioning at the host-pathogen interface, Incs are integral to mediating fusion with host-derived vesicles, forming membrane contact sites with host organelles, redirecting host vesicular transport to the inclusion, and modulating the host cell cytoskeleton^11–20^ all of which demands not only controlled interactions with host proteins but are also hypothesized to require interactions with other Inc proteins^21,22^. Given the retention of a large number of Inc proteins, it suggests that they play a critical role in the intracellular survival of *Chlamydia.* Indeed, recent work from our laboratory^23^ and others^19,20^ revealed that the absence of select Inc proteins, including CTL0481 (CT229 in serovar D), results in premature inclusion lysis and host cell death, leading to its designation as a <u>C</u>hlamydia <u>p</u>romoter <u>o</u>f <u>S</u>urvival (CpoS). While CpoS binds to and recruits multiple host Rab GTPases to control host trafficking pathways and suppresses cell-autonomous immunity^12,14,15,19,20,24^, CpoS also binds to other Inc proteins^19–21^. However, the significance of these Inc-Inc interactions remains poorly understood.

To acquire key nutrients from the host and membrane for the growing inclusion, *C.t.* modulates vesicular trafficking and fusogenicity with the inclusion membrane ^9,25^. Small guanosine triphosphate (GTP) binding proteins such as Rab GTPases and Arf GTPases regulate key aspects of vesicle trafficking, including vesicle formation, transport, tethering, and fusion^26,27^. Rab GTPases interact directly or indirectly with souble NSF (N-ethylmalemide-sensitive factor) attachment receptors (SNAREs)^28^ to regulate the assembly or disassembly of SNARE complexes^29,30^. In addition to hijacking small GTPases to modulate host vesicular transport, *C.t.* Inc proteins engage host SNARE proteins ^31–33^. Furthermore, bioinformatic analysis of Inc proteins has revealed that at least three of them—IncA, IPAM, and InaC—possess SNARE-like domains (SLD)^34^ that mediate homotypic inclusion fusion^16,33,35,36^. In eukaryotes, SNARE domains form heteromultimers with other SNARE domains. Typically, this involves one R-SNARE associating with three Q SNAREs, across the donor and target membrane, in addition to Rab GTPases and tethering factors^30,37,38^. Upon vesicle docking, a stable four-helix bundle is formed from the four SNARE domains aligning, allowing the zippering of the inner and outer leaflets of the donor and target membranes, leading to membrane fusion. A crucial feature of the SNARE domain is its highly conserved central ionic “0” layer, containing either a conserved arginine (R-SNARE) or glutamine (Q-SNARE). The salt bridges formed by the ionic layer, combined with hydrophobic interactions from surrounding leucine (heptad) repeats, initiate the zippering process that drives membrane fusion.

While the engagement of SNARE proteins and small GTPases is critical for modulating vesicular trafficking during *C.t.* infection, interactions among inclusion membrane proteins may play an equally vital role in promoting vesicle tethering and fusion. Advances in chlamydial genetics, along with new tools to capture dynamic protein-protein interactions, suggest that Inc proteins can bind to one another throughout the developmental cycle^19,20,22^. Notably, CpoS forms higher oligomeric structures, such as tetramers, which we posit enables it to simultaneously bind to both other Incs and host factors. Collectively, our work reveals that Inc-Inc interactions add to the complex organization of the inclusion membrane and collaboration between these bacterial proteins may enable the pathogen to more efficiently subvert host regulations. Significantly, we show that disruption of these interactions can perturb host-pathogen interactions compromising the integrity of the inclusion membrane.

## Materials and Methods

### Bacteria and cell culture

*Chlamydia trachomatis* serovar L2 (LGV 434/Bu) was propagated in HeLa 229 cells (American Type Culture Collection) grown in RPMI 1640 with L-glutamine (ThermoFisher Cat#11875-093) supplemented with 10% Fetal Bovine Serum (FBS) (VWR Cat#89510-186), 1mM sodium pyruvate (ThermoFisher Cat#11360070), and sodium bicarbonate. Cells were grown at 37°C in a humidified incubator with 5% CO_2_. When required, EBs were purified from HeLa cells using a gastrografin density gradient^39^.

### Cloning

For expression in *C.t.*, CpoS and various CC-containing Incs were PCR amplified from purified *C.t.* L2 genomic DNA, and a C-terminal HA-tag or FLAG-tag was added. The resulting PCR fragments were digested with KpnI-HF (NEB Cat# R3142L) and SalI-HF (NEB Cat# R3138L) and cloned into pBomb4. To truncate or specifically delete the CC domains, the regions determined using SMART sequence analysis, were removed using C-terminal truncations or internal deletions conducted by GenScript. For in vitro biochemical characterization, full-length (FL) or Inc proteins lacking the transmembrane domain were cloned into the pMALc-5VT vector (Protein and Crystallography Facility, UIOWA) using NotI (NEB Cat#R3189L) and SalI-HF, resulting in an N-terminal fusion to an MBP-tag and a C-terminal 6X-His-tag. Truncated CpoS was cloned into pGEX6P1 (Sigma Cat#GE28-9546-48) using BamHI-HF (NEB Cat#R3136L) and Xho1 (NEB Cat#R0146L) to generate an N-terminal fusion to a GST-tag. All primers are listed in Table S1. The integrity of all constructs was confirmed by sequencing (McLab).

### Immunoprecipitation

HeLa cells were infected at an MOI of 3 with *C.t.* L2 pBomb4-tet-CpoS HA-tag and one of the various pBomb4-tet-Inc FLAG-tag strains. Alternatively, HeLa cells were transfected with pcDNA3.1eGFP-Rab14 (Genscript Cat#OHu12700) or pcMV eGFP Rab35 (Addgene Cat#49552) followed by infection at an MOI of 3 with *C.t.* L2 pBomb4-tet-CpoS HA-tag and one of the various pBomb4-tet-Inc FLAG-tag strains. Expression of both *C.t.* proteins was induced with 10 ng/ml anhydrous tetracycline (aTc) at the time of infection. At 24 hpi, cells were lysed on ice in eukaryotic lysis solution (ELS) (50 mM Tris HCl, pH 7.4, 150 mM NaCl, 1 mM EDTA, and 1% Triton-X 100) and spun at 12,000 *× g* for 20 min. Supernatants were incubated with HA beads (anti-HA Magnetic beads, Thermo Fisher Scientific Cat#88836) for 1-2 h at 4°C. For CpoS truncations, FLAG beads were used for IP. The beads were washed 5 times with ELS without Triton-X 100. Proteins were eluted from the beads in NuPAGE LDS Sample Buffer (Thermo Fisher Scientific Cat#NP0007) by heating at 100°C for 5 min prior to analysis by western blotting.

### Western blotting

Samples were separated by sodium dodecyl sulfate-polyacrylamide gel electrophoresis (SDS-PAGE) on a 4-12% Bis-Tris protein gel (GenScript Cat#M00653) with MES running buffer. Proteins were transferred onto PVDF membranes and blocked in 5% milk in Tris-buffered saline with Tween 20 overnight at 4°C. Membranes were probed with anti-FLAG (Thermo Fisher Scientific Cat#701629) or anti-HA (1:1500 Millipore Sigma Cat#SAB2702217) primary antibodies and goat anti-rabbit HRP conjugate (1:10,000 BioRad Cat#1706515) secondary antibody. Results were collected from at least three independent experiments.

### Super-resolution microscopy

HeLa cells, seeded on #1.5 High Precision Glass Coverslips (Bioscience Tools Cat#CSHP-No1.5-13), were infected at an MOI of 5 with *C.t.* L2 pBomb4-tet-CpoS HA-tag and one of the various pBomb4-tet-Inc FLAG-tag strains. Expression of both proteins was induced with 10 ng/ml aTc at the time of infection. At 24 hpi, cells were fixed in 2% formaldehyde and permeabilized with 0.1% Triton-X 100. Samples were blocked with 3% BSA in phosphate-buffered saline (PBS) followed by a 1-hour incubation with primary antibodies, rabbit anti-HA (1:1000, Novus Cat#NB600-363) and mouse anti-FLAG (1:1000, Sigma Cat#F2555), prepared in blocking buffer. Cells were washed with PBS followed by incubation with the secondary antibodies goat anti-mouse DyLight594 (1:250, Invitrogen Cat#35561) and goat anti-rabbit STAR635 (1:250, Abberior Cat#ST635P-1002) prepared in blocking buffer. Coverslips were washed with PBS and mounted on glass slides using ProLong Diamond Antifade Mountant (Invitrogen Cat#36961) and allowed to cure for 24 hrs before imaging. STED images were taken using a Leica SP8 inverted microscope. Images were deconvoluted using Imaris Professional Software. Results were collected from at least three independent experiments.

### Confocal microscopy

HeLa cells, seeded on coverslips, were transfected with GFP-tagged Rab (Addgene #129020, #49467) and Arf GTPases (Addgene #49578, #39556) using Lipofectamine LTX. At 24 hrs post-transfection, cells were infected with *C.t.* L2 WT, *cpoS*::*bla*,^12,23^ *CT228:bla,* or *InaC::bla* at an MOI of 2. At 24 hpi, cells were fixed with methanol and blocked with 3% BSA in PBS. The inclusion membrane was stained with either anti-IncB or anti-IncA antibodies (1:250) diluted in blocking buffer. Cells were washed with PBS and incubated with goat anti-rabbit AlexaFluor594 secondary antibody (1:1000, Invitrogen) and DAPI (1:1000, Invitrogen Cat#D1306) in blocking buffer. After washing with PBS, cells were mounted on glass slides using ProLong Diamond Antifade Mountant and allowed to cure for 24 hrs before imaging. Confocal images were taken using a Nikon inverted microscope. Results were obtained from at least three independent experiments.

### Protein expression

Sequence-confirmed plasmids were transformed into BL21 (DE3) or Rosetta (DE3) cells, grown at 37°C to an OD600 of 0.6, induced with 1 mM isopropyl β-D-1-thiogalactopyranoside (IPTG) (Research Products International Cat#I56000-25.0), and incubated overnight at 18°C with shaking. For purification of MBP-tagged proteins, cells were pelleted and resuspended in sonication buffer (20 mM Tris-Cl, pH 7.5, 200 mM NaCl, 1 mM DTT, 1 mM EDTA, and 10% glycerol) containing a protease inhibitor cocktail (Millipore Sigma Cat#1187350001) and DNase I (Millipore Sigma Cat#10104159001). Cells were lysed by sonication (Sonics Vibram-Cell) at 38% amplitude with 1 s on/1 s off pulses. The soluble fraction was collected by high-speed centrifugation at 12,000 *x g* for 1 hr and applied to an Amylose resin high-flow column (NEB Cat#E8022S). Unbound proteins were removed by washing with sonication buffer, and bound proteins were eluted with sonication buffer containing 10 mM maltose (Research Products International Cat#M22000).

For purification of GST-CpoS C-ter, cells were lysed in lysis buffer (50 mM Tris-Cl, pH 7.5, 500 mM NaCl, 10% glycerol, 1 mM DTT, 1X PIC, and 5 µL of 50 units of DNase). The supernatant was applied to glutathione agarose beads (Thermo Fisher Scientific Cat#16100), washed with wash buffer (50 mM Tris-Cl, pH 7.5, 300 mM NaCl, 10% glycerol, 1 mM DTT), and eluted with elution buffer (50 mM Tris-Cl, pH 7.5, 150 mM NaCl, 10% glycerol, 10 mM reduced glutathione). Protein samples were resolved using 4-12% Bis-Tris SDS-PAGE gel and stained with Coomassie Brilliant Blue (Bio-Rad Cat#1610436). Protein identity was confirmed by western blotting using anti-GST (1:5000, Thermo Fisher Scientific Cat#MA4-004) or anti-MBP (1:5000, Santa Cruz Biotechnology Cat#sc-13564) antibodies. Proteins were further purified using a HiLoad 16/600 24 ml Analytical Superdex™ 200 Increase pg gel-filtration column (GE Healthcare) with 20 mM Tris, pH 7.5, 200 mM NaCl, 10% glycerol, and 1 mM DTT. Injection volume was 500 µL with aflow rate of 0.5 ml/min, and fractions were collected in 500 µL volumes. Purified protein was concentrated using Amicon ultra-centrifugal filter devices (Millipore Sigma Cat#UFC801024, 10 kDa cutoff) and assessed again using SDS-PAGE gel.

### *In vitro* binding assays using GST-pulldown

To assess *in vitro* binding, 500 µL of glutathione agarose resin was equilibrated with 10 ml of column buffer (50 mM Tris, pH 7.4, 500 mM NaCl, 10% glycerol, 1 mM DTT). Twenty micrograms of GST-tagged CpoS C-ter were added and incubated for 2 hrs at 4°C. After three washes with column buffer, 20 µg of MBP-tagged Incs (MBP-InaC C-ter, MBP-IPAM C-ter, MBP-CT449, or MBP control) were added and incubated overnight at 4°C with rotation. The column was washed three times with wash buffer to remove unbound proteins. The protein complex was eluted by incubating the resin with a GST elution buffer for 30 min. The flow-through was collected, and the complexes were confirmed by western blotting with anti-GST and anti-MBP antibodies and goat anti-mouse HRP conjugate secondary antibody. Results were obtained from at least three independent experiments.

### Mass photometry

To determine the oligomeric states of CpoS in solution and accurately measure molecular mass, mass photometry (MP) was performed using a Refyn TwoMP mass photometer (Refeyn Ltd, Oxford, UK). Microscope coverslips (24 mm x 50 mm, Thorlabs Inc.) and silicon gaskets (Grace Bio-Labs) were cleaned by serial rinsing with Milli-Q water and HPLC-grade isopropanol (Sigma Aldrich Cat#34863) followed by drying with a filtered air stream. All MP measurements were performed at room temperature using Dulbecco’s phosphate-buffered saline (DPBS) without calcium and magnesium (Gibco Cat#14200-075). The instrument was calibrated using a protein standard mixture: β-amylase (Sigma-Aldrich Cat#A7005-10KU, 56, 112, and 224 kDa) and thyroglobulin (Sigma-Aldrich Cat#T1001-100MG, 670 kDa). In brief, 15 µL of DPBS buffer was placed in the well to find focus before each measurement. The focus position was locked using the default droplet-dilution autofocus function, after which 5 µL of protein (100 nM) was added and briefly mixed before movie acquisition started. Movies were acquired for 60 s (3000 frames) using AcquireMP (Refeyn Ltd) under standard settings, and the contrast values were converted to molecular masses using the instrument’s calibration function. After data collection, the coverslip was repositioned to place the next channel or sample well over the objective for consecutive measurements. MP data were plotted as histograms or kernel density estimate (KDE) distributions. The distribution peaks were fitted with Gaussian functions to obtain the average molecular mass of each distribution component. All movies were processed and analyzed using DiscoverMP (Refeyn Ltd).

### Size Exclusion Chromatography Coupled with Multi-Angle Light Scattering and Small Angle X-ray Scattering (SEC-MALS-SAXS)

For detailed biophysical analysis of CpoS proteins in solution, SEC-MALS-SAXS was performed. Datasets were collected using the 18-ID-D BioCAT Beamline at the Advanced Photon Source (APS) at Argonne National Laboratory (Chicago, IL). Samples were centrifuged for 5 min at 13,000 rpm to remove any potential aggregates prior to column loading. Samples containing 1 mg/ml of MBP-tagged CpoS in 250 µL were injected onto a 24 ml Superdex 200 Increase 10/300 GL column (GE) equilibrated with buffer (20 mM Tris, pH 7.5, 50 mM NaCl, 1 mM EDTA, 2% glycerol, and 1 mM DTT) at a flow rate of 0.6 ml/min on an Agilent 1300 chromatography system. Column eluant was analyzed in line by the UV absorbance detector of the Agilent 1300 chromatography system, then directed into the DAWN Heleos-II light scattering (LS) and OptiLab T-rEX refractive index detectors in series. Finally, the elution trajectory directed samples into a 1.0-mm ID quartz capillary SAXS sample cell. Scattering data were collected every 1 second using a 0.3-second exposure and detected with an Eiger2 XE 9M pixel detector (DECTRIS) with a 12 KeV (1.033 Å wavelength) X-ray beam covering a q-range of 0.0027 < q < 0.41 Å-1 (q = 4π*sinθ/λ, where λ is the wavelength and 2θ is the scattering angle). Accurate protein molecular weights from MALS data were determined using the ASTRA software (Wyatt Technology).

### Small-angle X-ray scattering (SAXS) data processing and modeling

SAXS data reduction, buffer subtraction, and further analysis were performed using BioXTAS RAW version 2.1.4^40^. An average of 30 frames before and after the eluted peaks were used for buffer subtraction. Protein peaks were also ran through evolving factor analysis (EFA) to deconvolute peaks into the individual scattering components where applicable^41^. The forward scattering intensity I (0) and radius of gyration (Rg) were calculated from the Guinier fit. The normalized Kratky plot and pair distance distribution P(r) plot were calculated using the program GNOM embedded in the BioXTAS RAW software^42^.

### Statistical Analyses and Densitometry

When necessary, statistical analysis was performed using GraphPad Prism 9.3.0 software. One-way ANOVAs were used followed by Tukey’s multiple comparisons with *P* < 0.05 (*), *P* < 0.01 (**), and *P* < 0.001 (***).

## Results

### CpoS binds to multiple Inc proteins

CpoS was previously confirmed to bind to IPAM during infection^19^. However, we hypothesized that it may engage other Inc proteins throughout the chlamydial lifecycle. As coiled-coil (CC) domains are found in numerous proteins and mediate important protein-protein interactions (PPI) that control diverse cellular processes, we sought to determine whether CpoS might bind to CC-domain-containing Incs. Using the Simple Modular Architecture Research Tool (SMART), we evaluated the 37 Inc proteins previously shown to localize to the inclusion membrane^43^. In line with previous reports^34^, we identified nine Inc proteins that possess a CC domain: IncA, CTL0476/CT222, CTL0476/CT223 (IPAM), CTL0477/CT224 (Tri1), CTL0478/CT226, CTL0480/CT228, CTL0485/CT233 (IncC), CTL0540/CT288 (IncM), and InaC (Figure 1A).

**Figure 1:**
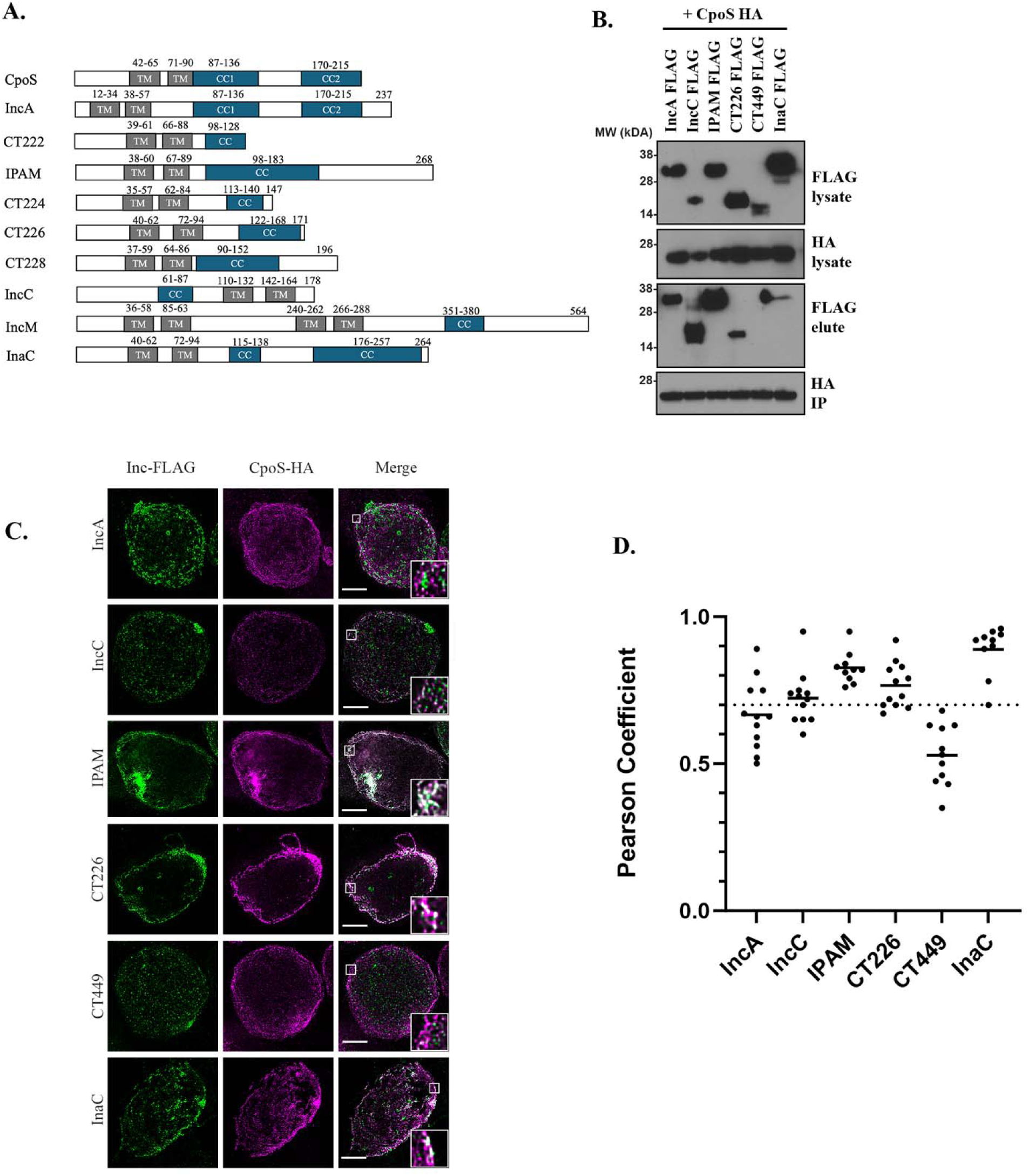
CpoS binds to multiple coiled-coil domain containing Incs. (**A)** Schematic of Inc proteins. Bi-lobed hydrophobic domain is shown in gray and the coiled-coil domain is represented in teal. (**B, C**) HeLa cells were coinfected at an MOI of 2.5 with HA-tagged CpoS and FLAG-tagged Inc for 24hrs. (**B**) CpoS was immunoprecipitated from cell lysates using HA-magnetic beads and samples were analyzed by western blotting. (**C)** Cells were formaldehyde fixed 24hpi and processed for super-resolution microscopy using anti-HA and anti-FLAG antibodies with anti-mouse 594 and anti-rabbit 635 secondaries, respectively. Scale bar denotes 5μM. White boxes denote the area used for inset. Pearson Correlation Coefficients (R Value) were calculated using ImageJ Coloc2 function. The graph is representative of 10 images per experiment. Dashed line represents cutoff for significant colocalization (0.70). (**B, C**) Data are representative of 3 independent experiments.

Due to a lack of commercially available Inc antibodies, we used a co-infection model in which HeLa cells were infected with two strains: *C.t.* CpoS-HA and another Inc fused to a FLAG-tag. Since inclusions fuse, this model results in a single inclusion per cell with both Incs present on the same inclusion membrane^19^. Using anti-HA magnetic beads, we enriched for the CpoS and probed immunoblots with anti-HA to confirm the IP and anti-FLAG to detect interactions with CC-domain containing Incs. Among the Inc proteins evaluated for their ability to bind CpoS, we detected binding with IncA, IncC, IPAM, CT226, and InaC (Figure 1B). The identified Inc proteins exhibited consistent interactions with CpoS, with the exception of IncA, which failed to bind in multiple attempts. This suggests that IncA may have a weaker affinity for CpoS compared to the other Inc proteins or interact transiently. Due to low expression of CT222, Tri1, CT228, and IncM we could not accurately ascertain whether they similarly bound to CpoS and thus they were excluded from subsequently analysis.

We next sought to confirm that CpoS co-localizes with these Inc proteins. Using STED super-resolution microscopy to acquire high-resolution images of the inclusion membrane, we demonstrate that CpoS co-localizes with the CC-domain-containing Incs IncC, IPAM, CT226, and InaC (Figure 1C), and to a lesser extent with IncA, which exhibited a lower colocalization score. Importantly, CpoS did not colocalize with CT449, an Inc protein that lacks coiled coil domains and does not bind CpoS (Figure 1B). Overexpression of CpoS did not alter the localization of any of the tested Incs. Taken together, these results suggest that CpoS interacts with multiple CC-domain-containing Incs during *C.t.* infection.

### CpoS interacts with the C-terminal region of Incs

Given the crucial role of CC domains in mediating protein-protein interactions (PPIs), we hypothesized that this domain is necessary for binding to CpoS. To test this, we created truncation constructs of two Inc proteins, IPAM and InaC), resulting in versions either lacking only the C-terminus or the CC domain (Figure 2A). For InaC, which contains two CC domains, two constructs were generated: one lacked only CC1 (inner), while the other lacked CC2 (outer) (Figure 2A). Cells were co-infected with full-length CpoS-HA and the *C.t.* strain expressing the truncated or CC-deletion FLAG-tagged Incs. As shown in Figure 2B, deletion of the C-terminus, but not the CC domain from IPAM, resulted in the loss of CpoS binding. In contrast, for InaC, deletion of CC2 led to a loss of binding to CpoS, while CC1 was dispensable for the interaction. Collectively, our results indicate that CpoS binding to CC domain-containing Incs is context-dependent, requiring specific structural features, with different regions contributing variably to the interaction.

**Figure 2:**
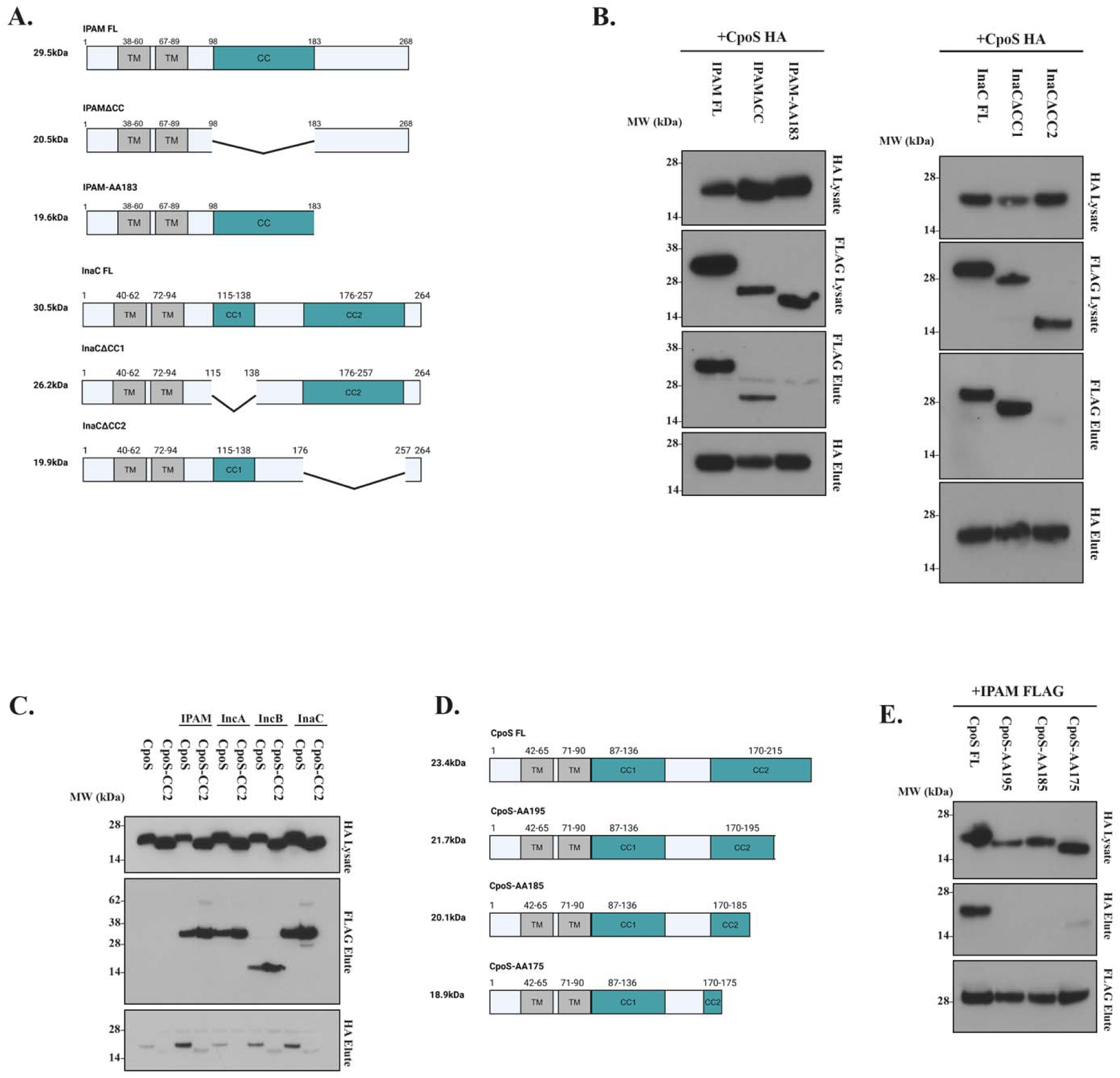
Outer regions of Incs are involved in Inc-Inc interaction. **(A)** Schematic of Inc truncations. Trunctations were made to remove large regions of predicted functional domains. **(B)**) HeLa cells were coinfected at an MOI of 2.5 with HA-tagged CpoS and FLAG-tagged Inc for 24hrs. CpoS was immunoprecipitated from cell lysates using HA-magnetic beads and samples were analyzed by western blotting. **(C)** HeLa cells were coinfected at an MOI of 2.5 with either HA-CpoS FL or HA-CpoS-CC2 and FLAG-tagged Inc for 24hrs. The tagged Incs were immunoprecipitated from cell lysates using FLAG-magnetic beads and samples were analyzed by western blotting. **(D)** Schematic of sequential truncations of CC2 in CpoS. **(E)** HeLa cells were coinfected at an MOI of 2.5 with the HA-tagged CC2 truncations and FLAG-tagged IPAM for 24hrs. IPAM was immunoprecipitated from cell lysates using FLAG-magentic beads and samples were analyzed by western blotting. (**B, C, E**) Data are representative of 3 independent experiments.

### The CC2 domain of CpoS is crucial for mediating Inc-Inc interaction

Previous studies demonstrated that CpoS interacts with IPAM through its CC2 domain^19^. Building on these findings, we investigated the necessity of the CC2 domain interactions with other Inc proteinss. We generated CpoS constructs lacking either CC1 and CC2 and confirmed that while CC1 is essential for Rab binding, CC2 is dispensable (Figure S1). We then conducted co-infection immunoprecipitation assays using either full length (FL) CpoS or a CpoS construct lacking CC2, co-infected with FLAG-tagged IPAM, IncA, IncB, or InaC. Anti-FLAG beads were used to enrich the CC-Incs and assess their interaction with the different CpoS constructs. As anticipated, we observed a significant decrease in the retention of CpoS-ΔCC2 compared to FL CpoS for each Inc IP (Figure 2C), indicating that the CC2 domain plays a vital role in facilitating interactions with multiple Inc proteins.

Next we sought to pinpoint the specific region within CC2 required for these interactions. Sequential 10-amino-acid truncations of the CC2 domain were generated (Figure 2D), and co-infection IP assays were repeated to identify truncations that disrupted binding. Our results indicate that the last 20 amino acids of CC2 are required for binding (Figure 2E). Taken together, these results demonstrate that the outer region of CpoS, IPAM, and InaC is involved in Inc-Inc interactions, while the internal domains may be dedicated to interactions with host proteins.

### CpoS binds to Rabs and Incs simultaneously

CpoS contains two coiled-coil domains, with CC1 implicated in binding to Rab GTPases and CC2 mediating interactions with multiple Inc proteins (Figure 2)^19^. This dual-domain structure suggests that CpoS may serve as a molecular hub coordinating interactions between host Rab GTPases and other bacterial Inc proteins. To test this hypothesis, we transfected HeLa cells with GFP-tagged Rab14 and Rab35 and used our Inc co-infection model to evaluate Inc and Rab binding to CpoS. Immunoprecipitation of CpoS-HA revealed that it can simulanteously associate with both Rab GTPases and other Inc proteins (Figure 3). These findings suggest a central role for CpoS in linking host and bacterial factors, potentially facilitating the dynamic regulation of inclusion membrane function during infection.

**Figure 3:**
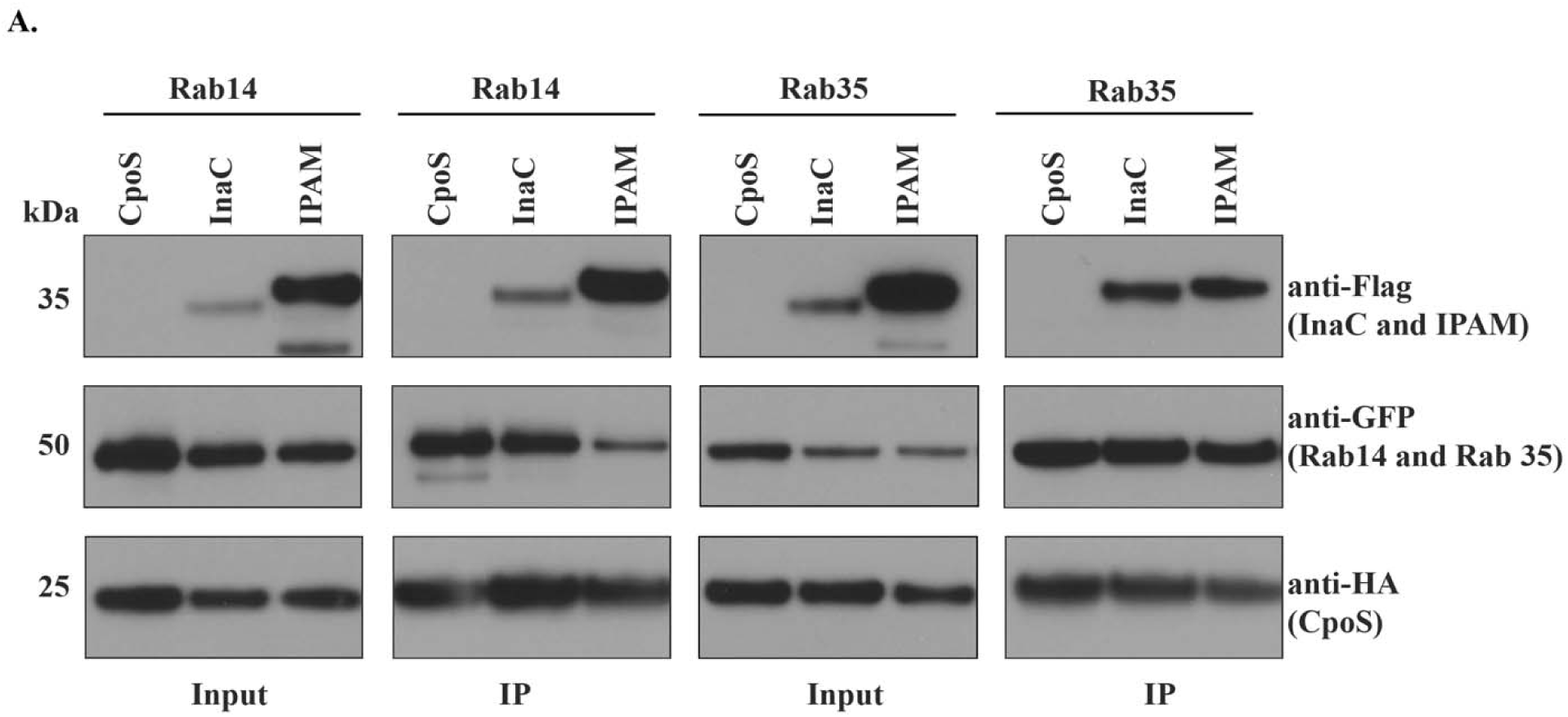
CpoS simultaneously interacts with Rab GTPases and other *C.t* Incs. (A) HeLa Cells were transfected with pcDNA 3.1 eGFP-Rab14 or pcMV eGFP Rab 35 and co-infected at an MOI of 5 with *C.t* expressing CpoS-HA and either *C.t* expressing IPAM-Flag or InaC-Flag. After 24hpi, CpoS was immunoprecipitated using HA beads and the elutes were resolved on SDS-PAGE followed by western blotting with HA, GFP and Flag antibodies.

### Tetramer formation by CpoS requires CC2

Currently, the molecular architecture of CpoS is poorly understood, limiting our ability to understand the role it plays in modulating both other Chlamydial Incs and Rab GTPases. To address this, we conducted biochemical and biophysical analysis of the protein. CpoS, lacking the N-terminus and the bilobed hydrophobic domain (CpoS C-ter) (Figur 4A) was expressed as an MBP-fusion in *E. coli.* Protein was isolated from bacterial lysates using affinity chromatography and was further purified using Superdex™ 200 at a flow rate of 0.5 ml/minute. Coomassie staining and western blotting of the eluants confirmed highly pure protein was obtained (Figure S2A). Intriguingly a higher molecular weight SDS resistant species of CpoS C-ter was noted by western blotting (Figure S2A, asterick), suggesting it might oligomerize. Size-exclusion chromatography (SEC) of CpoS C-ter yielded three well-separated peaks, further suggesting CpoS forms higher-ordered structures (Figure 4B). To rule out the possibility of protein aggregation, dynamic light scattering (DLS) and mass photometry (MP) (Figure 4B) was used to evaluate the nature of the proteins present within each peak. As shown in Figure 4B, CpoS C-ter predominately formed tetramers (∼242 kDa) and octomers (∼488 kDa) in solution along with some MBP-tag and monomeric CpoS C-ter (∼58 kDa). CpoS C-ter also formed distinct monomers, tetramers and octomers in some of the fractions. Notably, CpoS oligomerization was pH dependent, with the tetramer being stable at neutral pH and dissociating into dimers and monomers in both acidic and basic buffer conditions (Figure S2C).

**Figure 4:**
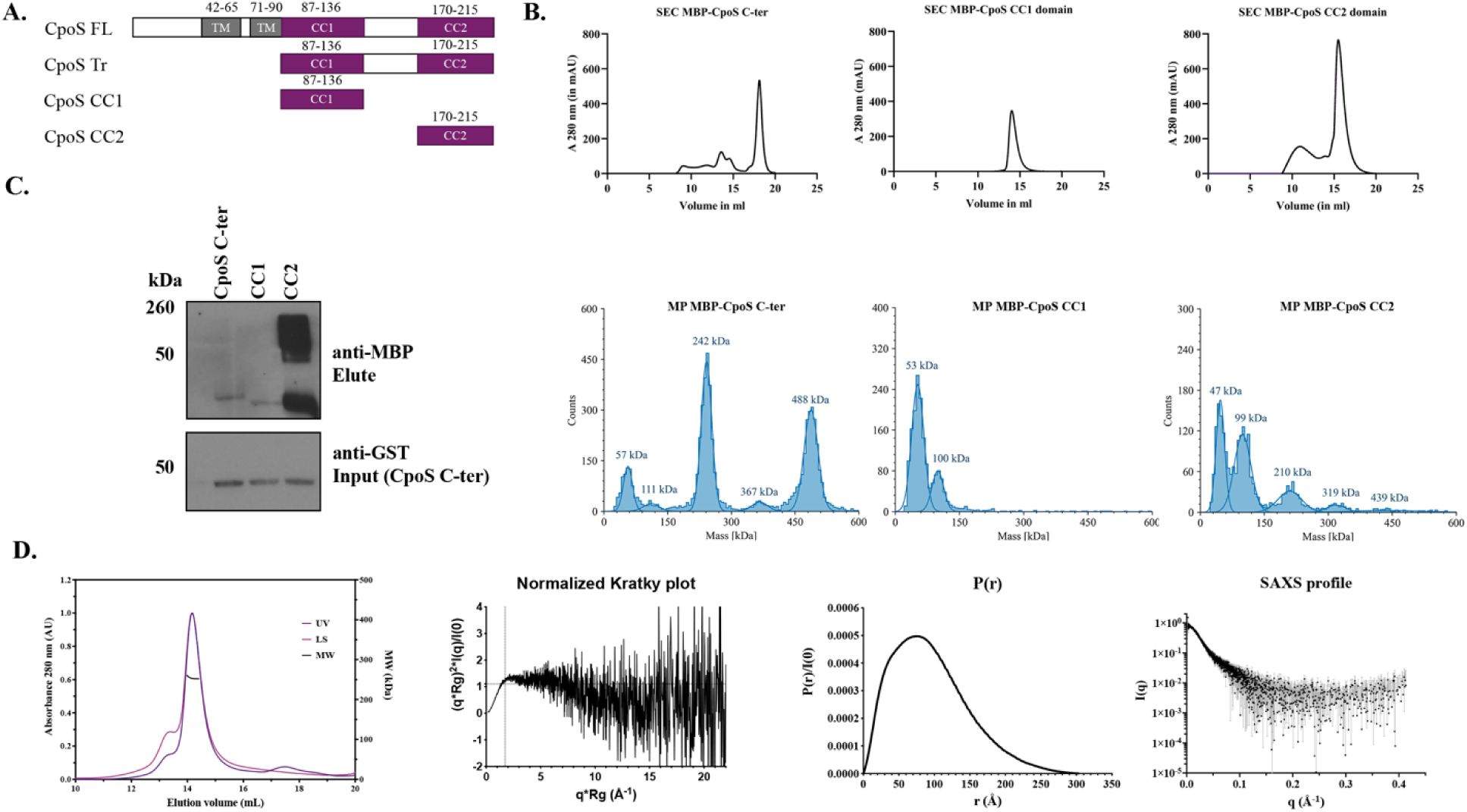
Tetramerization property of CpoS C-ter. (A) Domain organization of various truncations of CpoS protein used in the study. MBP-tagged fusion proteins of CpoS C-ter, CC1 domain and CC2 domain were expressed in (BL21 DE3) cells for all the invitro biophysical studies. (B) SEC chromatogram of 500 µl (injection volume) of CpoS C-ter on AS 200 column with a flow rate of 0.5 ml/minute. Mass photometry histograms of CpoS C-ter (eluted from SEC column) confirmed the dominating tetramers and octamers of the CpoS C-ter proteins in solution and oligomerization of CC2 domain (C) GST-pulldown assays analyzed by western blotting to show molecular interaction of GST-CpoS C-ter with MBP-CpoS C-ter, confirming the CC2 domain was sufficient for tetramerization. (D) SEC-MALS profile of CpoS C-ter tetramer on S-200 column. Kratky plot showing the folding nature of CpoS C-ter. PDDF fitting curves derived from the SAXS profile. SAXS profile for CpoS C-ter. The plot represents the logarithm of the scattering intensity (I, artibrary units) as a function of momentum transfer (s, in Å^−1^).

To determine whether assembly and oligomerization requires CC1 or CC2 of CpoS, we expressed each domain independently and analyzed purified protein samples by SEC and MP (Figure S2A, 4B). Similar to what was observed for CpoS C-ter, a higher molecular weight species was noted by western blotting for CC2 but not CC1 (Figure S2A, asterick). The requirement of the CC2 domain for tetramer and octamer formation was further confirmed by SEC, MP and native PAGE, (Figure 4B, 5B), revealing only CpoS CC2 was capable of oligomerizing. We hypothesized that the formation of a CpoS tetramer required interactions between its CC2 domain. To test this, we conducted *in vitro* pulldowns using GST-CpoS C-ter with MBP CpoS C-ter, MBP-CC1, MBP-CC2 domains. As predicted, CC2 was required for interaction with itself and tetramerization (Figure 4C).

To further characterize the tetrameric structure of CpoS C-ter, size exclusion chromatography coupled with multiangle light scattering and small angle X-ray scattering (SEC-MALS-SAXS) was employed. The theoretical mass of the MBP-tagged Cpos C-ter tetramer is 234 kDa, as the molecular weight of the monomer, derived from the protein sequence, is 58kDa. The absolute molecular mass of CpoS C-ter, measured by MALS was 254 kDa and 263 kDa as measured by SEC-SAXS (Figure 4D). These data are consistent with tetrameric assembly of CpoS C-ter.

Following data reduction and buffer subtraction, the SAXS data were further processed using the BioXTAS RAW software ^40^. The forward scattering intensity I(0) and the radius of gyration (Rg) were calculated from the Guinier fit. The normalized Kratky plot, the pair distance distribution plot P(r) and the Porod volume were calculated using GNOM embedded in BioXTAS RAW^44^. Analysis of the SAXS data indicated that the CpoS C-ter protein was monodisperse, as evidenced by the linearity of the Guinier plot (Figure 4D). The normalized Kratky blot exhibited a maximum shift towards higher qRg values, and the P(r) function showed a peak at low radii followed by an extended tail, both suggesting an elongated shape for CpoS C-ter (Figure 4D). The molecular weight of CpoS C-ter, estimated from SAXS data using various methods was consistent with a tetrameric assembly: 262.8 kDa (by Vc method), 274.3 kDa (by Vp method), and 242.6 kDa (by Bayes method) ^45–47^. The Rg value derived from the Guinier approximation was 5.45 nm, while the P(r) analysis yielded an Rg of 5.47 nm and a maximum particle dimension (Dmax) of 19.5 nm. These parameters further support the elongated nature of the CpoS C-ter tetramer in solution. In summary, our SEC-MALS-SAXS analysis provides strong evidence for a stable, elongated tetrameric conformation of CpoS C-ter in solution, corroborating our previous biophysical characterizations (Figure 4D).

### Other Inc proteins undergo oligomerization

To investigate whether other CC domain-containing Inc proteins oligomerize, the purified proteins were analyzed by SEC, MP, and native PAGE. IncA, previously shown form tetrameric structures, was included as a control^36^. Our analysis revealed that IPAM C-ter and InaC C-ter formed dimers and trimers, while IncA C-ter formed a stable tetramer (252 kDa based on sequence), similar to the one seen for CpoS C-ter (234kDa based on sequence). As expected, CT449, which lacks a CC domain, exclusively formed monomers (Figure 5A). Oligomerization was further confirmed using native PAGE gel (Figure 5B).

**Figure 5:**
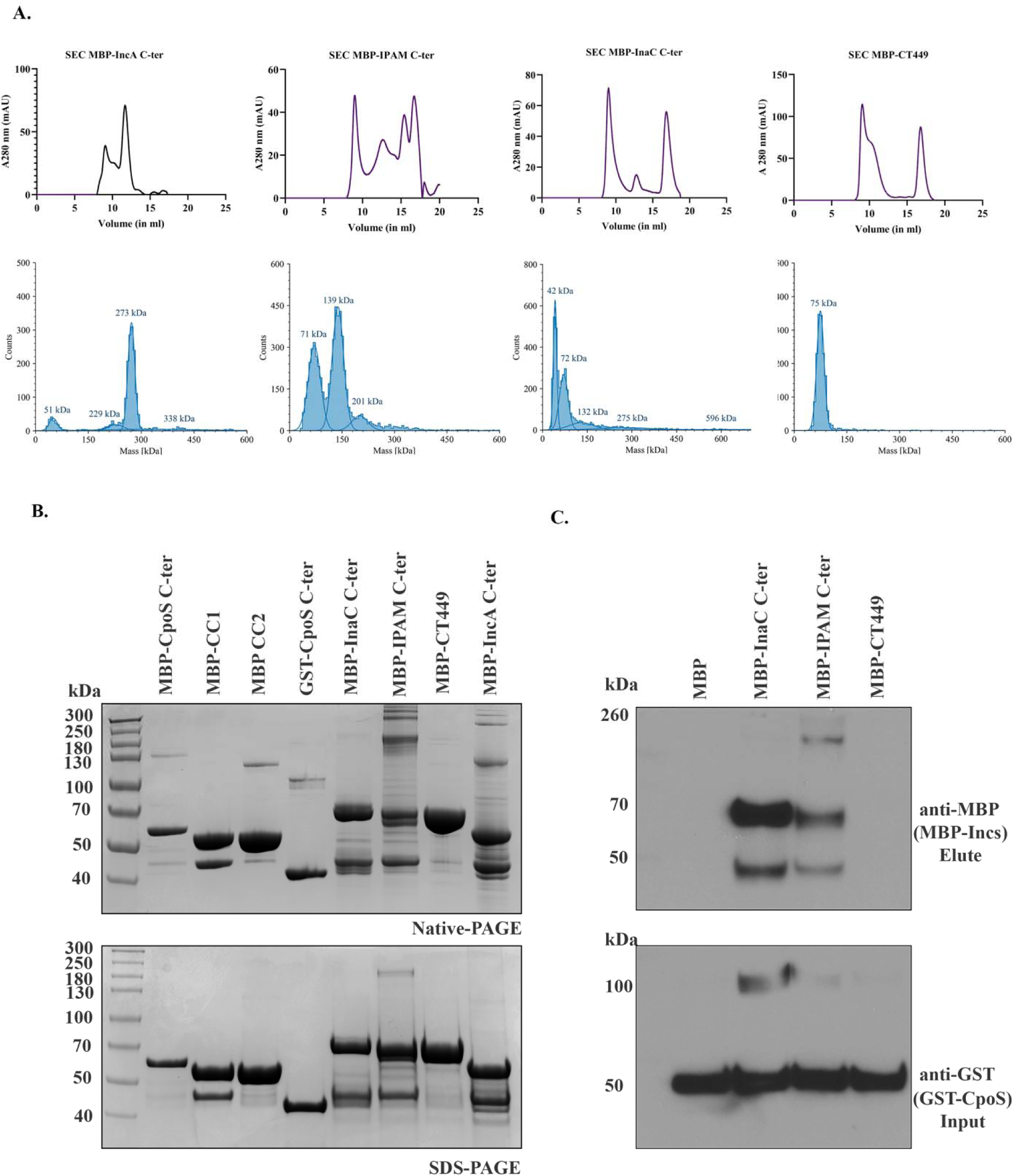
Intermolecular interactions of CpoS C-ter with other *C.t* Incs. (A) SEC chromatogram and MP to confirm the oligomerization property of other Chlamydial Inc proteins. The SEC profiles for IncA C-ter, IPAM C-ter, InaC C-ter eluted as multiple peaks which gave hints about possible oligomers for other Incs and not CT449 proteins, for which the proteins eluted as single peak and a void peak. The respective MP histograms for each Incs confirmed the presence of oligomers for IncA C-ter, IPAM C-ter, InaC C-ter, and monomers for CT449.

To confirm that CpoS binds directly to other CC domain-containing Inc proteins, we performed *in vitro* pull-down assays, using GST-CpoS. Notably, MBP-InaC and MBP-IPAM, but not MBP-CT449 or MBP tag alone, bound to CpoS (Figure 5C). These results are consistent with our IP’s (Figure 1B) and provide strong evidence that CpoS binds to multiple CC domain-containing Inc proteins.

### CpoS-InaC interactions are required for InaC-mediated recruitment of Arf GTPases to the inclusion membrane

Given the results showing multiple CC Incs bind to and colocalize with CpoS, we next sought to determine whether Inc-Inc interactions promote interactions with the cognate host factors. InaC is a well characterized Inc that interacts with Arf GTPases 1and 4^17,18^, host proteins involved in ER to Golgi trafficking^48^. To determine whether CpoS expression is required for InaC-mediated Arf1 or Arf4 recruitment, we assessed Arf1 recruitment to WT *C.t., cpoS::bla*, or *inaC::bla* inclusions. As shown in Figure 6A, Arf recruitment to the *cpoS::bla* mutant was impaired. To rule out this is not an artifact of premature inclusion lysis, only intact inclusions, as evident by IncB staining, were analyzed. Pulldown experiments confirmed that CpoS did not bind to Arf1 and Arf4, (Figure S3) further suggesting impaired recruitment is due to an inability to engage InaC. Further, to rule out a general growth defect causing the impaired recruitment, we repeated colocalization analysis using a CT228*::bla* strain, which did not show a significant decrease in colocalization (Figure 6A).

**Figure 6:**
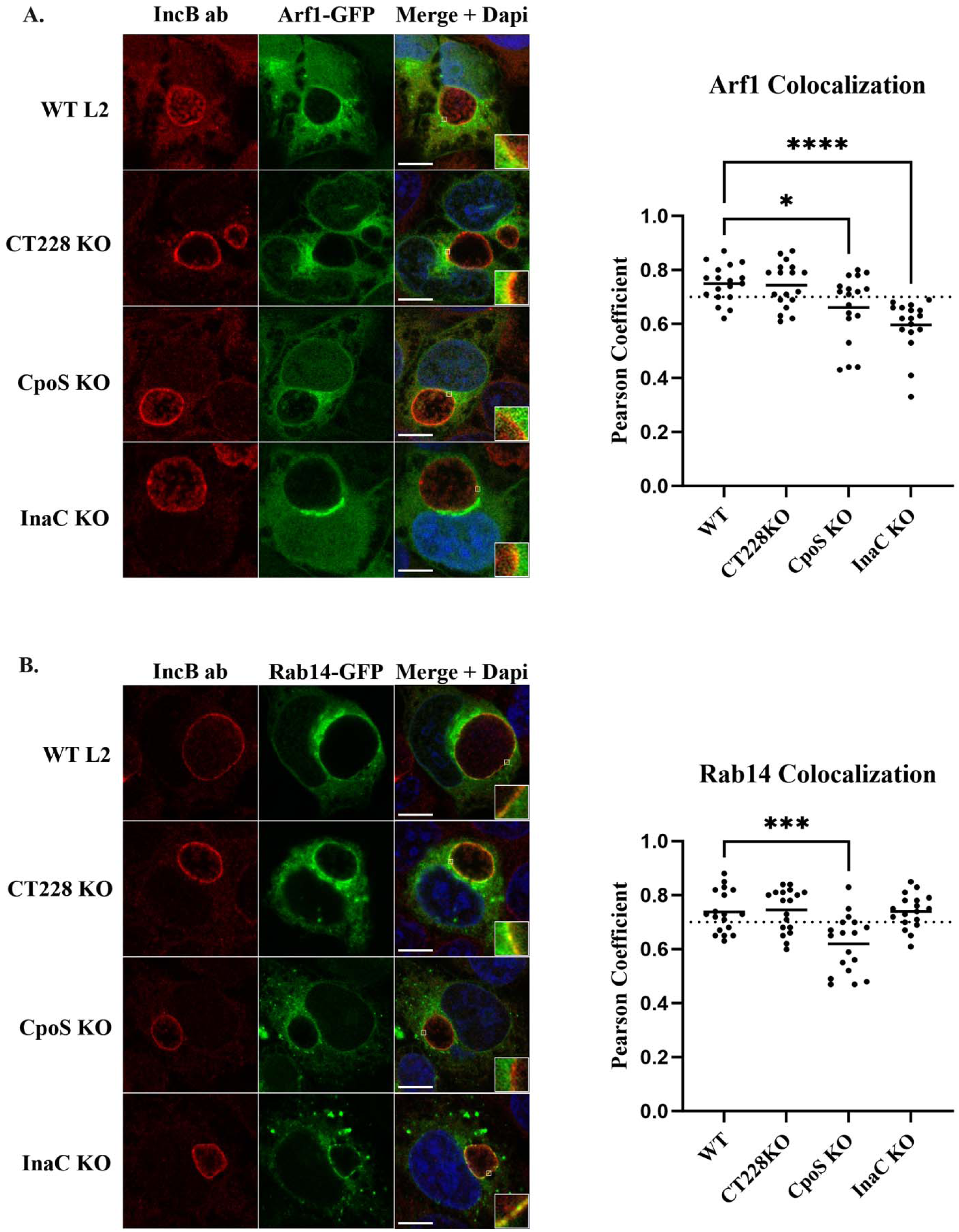
Loss of CpoS significantly affects Arf1 recruitment. HeLa cells were transfected with pEGFP Arf1 or pcDNA 3.1 Rab14 and infected with either WT L2, CT228*::bla*, *cpoS::bla*, or *inaC::bla C.t.* strains. Cells were MeOH fixed 24hpi and processed for confocal imaging. Intact inclusions were stained using an IncB specific antibody and with ant-rabbit 594 secondary and DAPI. Scale bar denotes 20μM. White boxes denote the area used for inset. Pearson Correlation Coefficients (R Value) were calculated using ImageJ Coloc2 function. The graph is representative of 18 images per experiment. Dashed line represents cutoff for significant colocalization (0.70). Data are representative of 3 independent experiments.

To determine whether Rab recruitment depends on CpoS binding to additional Inc proteins, we assessed Rab localization in cells infected with each of the mutants. Rab recruitment to the *inaC::bla* mutant showed no differences (Figure 6B), indicating that while CpoS-InaC interactions are critical for recruiting Arf GTPases, Rab GTPase recruitment occurs independently of Inc-Inc interactions. Similarly, testing the CT228*::bla* mutant revealed no significant changes in Rab colocalization. These results demonstrate that while CpoS-InaC interactions are essential for Arf protein recruitment to the inclusion, Rab GTPase recruitment relies on a distinct mechanism, highlighting a supportive role for Inc proteins in host protein localization.

## Discussion

In this study, we demonstrate that CpoS plays a multifaceted role during *C.t.* infection by simultaneously interacting with host Rab GTPases and various CC-domain-containing Inc proteins. Building on previous research, we show that CpoS specifically binds to itself and multiple other Inc proteins via its CC2 domain—a domain distinct from that required for Rab binding. Through these interactions, CpoS efficiently recruits diverse host vesicles to the inclusion, thereby supporting intracellular infection. Using biochemical and biophysical approaches, we show that CpoS forms higher-order structures, including tetramers and octomers, which we propose enables it to function as a scaffold for vesicle tethering and fusion—critical processes for delivering nutrient-rich cargo to the inclusion. The significance of this interactive network, centered around CpoS, is further highlighted by its essential role in maintaining the structural integrity of the inclusion membrane and evading the host immune response^19,20,23^.

The chlamydial inclusion serves as a protected intracellular niche that allows *C.t.* to replicate while evading host detection and the innate immune response. Genetic studies have highlighted the significance of several Incs in this process, revealing that the absence of select Incs leads to an unstable inclusion membrane that ultimately lyses, exposing the bacteria to the cytosol, and an inability to counter host defenses^20,23,49,50^. While the function(s) of a limited number of Incs are defined ^13,15,22,50–52^, CpoS has been shown to selectively recruit Rab GTPases to the inclusion membrane in a predominantly coiled-coil domain (CC1) dependent manner^12,19,20^, with some Rabs requiring additional residues in their C-terminus. This broad recruitment of Rab GTPases is essential for capturing lipid-containing vesicles and dampening STING activation and the interferon response ^19^. Notably, Rab35 recruitment is crucial for CpoS’s ability to moderate host responses; however, it also requires other unidentified factors. Furthermore, interactions between CpoS and Rab proteins appear to play a role in preventing premature cell death associated with CpoS mutant infections, suggesting that CpoS may have additional regulatory activities that influence this process.

*C.t.* is predicted to encode approximately 60 Incs, of which 37 have been confirmed to localize to the inclusion during infection^8,53,54^. Incs are strategically positioned on the cytoplasmic face of the inclusion, allowing them to interact with host factors while also engaging in self-interactions and interactions with one another. Early studies using bacterial two-hybrid assays and affinity purification mass spectrometry (AP-MS) have demonstrated that CpoS binds to various Inc proteins^21^, including confirmed interactions with IPAM during infection ^19,20^. We hypothesized that CpoS may also interact with other coiled-coil domain-containing Incs, and we successfully confirmed binding to CT226, InaC, CT228, and IncC. *C.t.* transcriptional activity can be broadly categorized into early, mid, and late stages, resulting in the secretion of effector proteins in waves throughout the developmental cycle^55^. Thus, while CpoS is expressed throughout this cycle, other Incs such as IPAM and InaC are primarily expressed during mid-cycle stages. This temporal expression pattern suggests that CpoS may interact with different partners at various stages of infection, potentially influencing the dynamics of Inc-Inc interactions and their functional outcomes.

Recent studies indicate that Inc-Inc interactions play a central role in organizing the inclusion membrane. Inclusion microdomains, defined regions within the inclusion, are thought to serve as molecular hubs that regulate the trafficking of inclusions to centrosomes^56,57^, maintain their stable association with the microtubule architecture^58^, and facilitate bacterial exit through extrusion ^13^. Previous research has shown that IncA and CpoS are essential for localizing microdomain Incs on the inclusion ^19,32^. CpoS has been shown to bind the microdomain Inc IPAM^19^, and our new data indicates it also binds the microdomain Inc, IncC. Due to the limited availability of Inc antibodies, we were unable to determine whether CpoS is similarly required for the localization of these other Incs to the inclusion membrane. Notably, both IncC and CpoS are important for bacterial replication and inclusion development as their absence leads to premature lysis. Prior studies have shown that different coiled-coil domains of CpoS have distinct functions in combating cell-autonomous defenses, with CC2 being critical for suppressing host cell death^19^. This raises the intriguing possibility that Inc-Inc interactions are crucial for forming an intact replicative niche, possibly through membrane acquisition, to support the expanding inclusion and facilitate the organization of microdomains.

Combining biochemical and biophysical methodologies to analyze the CC-domain containing Incs, we provide evidence that they are capable of forming higher ordered structures and some are capable of self-interactions. Their ability to oligomerize and interact with themselves and one another highlights the critical role Incs play in the formation of multi-protein complexes within the chlamydial inclusion membrane, which may enhance interactions with host proteins to modify host cell physiology. Early investigations using bacterial two-hybrid assays suggested that Inc proteins are capable of oligomerization, with specific emphasis on IncA, whose oligomerization requires its SNARE-like domain. Our biochemical studies further support this notion, revealing that multiple Incs, including CpoS, oligomerize to form higher-order structures and this also requires the C-terminal domain. We propose a model in which these proteins insert into the inclusion membrane via their transmembrane domains, while their coiled-coil tails form putative parallel four-helix bundles. This structural arrangement may facilitate membrane fusion events, akin to the mechanisms employed by eukaryotic SNARE proteins during vesicle docking and fusion. The parallels between Inc proteins and eukaryotic SNAREs are particularly noteworthy. Eukaryotic SNAREs are known to oligomerize into stable complexes that drive membrane fusion processes essential for intracellular transport. Our FoldSeq analysis confirmed that CpoS shares structural homology with t-SNARE proteins from *Trypanosoma cruzi* and v-SNARE proteins from *Arabidopsis thaliana*, despite minimal sequence similarity (Figure S4). This suggests that CpoS may perform similar functions in mediating membrane interactions and fusion processes within the chlamydial inclusion. The oligomerization of SNAREs is fundamental to their function, as it allows for the formation of a stable core complex that brings membranes into close proximity, facilitating fusion. Understanding how Inc proteins like CpoS utilize similar oligomerization strategies could provide valuable insights into the mechanisms by which *C.t.* manipulates host cellular processes for its benefit, highlighting a critical area for future exploration in host-pathogen interactions.

## Acknowledgements

We acknowledge grant support from the NIH (M.M.W., R01 AI150812, R01 AI155434, and R61 AI179999; XT T32 AI007511) and the University of Iowa Stead Family Scholars to M.M.W. We thank Tom Moninger from the University of Iowa Central Microscopy Research Facility for training in STED Microscopy.

